# Excitable dynamics through toxin-induced mRNA cleavage in bacteria

**DOI:** 10.1101/396226

**Authors:** Stefan Vet, Alexandra Vandervelde, Lendert Gelens

**Affiliations:** Applied Physics Research Group, Vrije Universiteit Brussel (VUB), Brussels, Belgium; Laboratory of Dynamics in Biological Systems, KU Leuven, Leuven, Belgium; Interuniversity Institute of Bioinformatics in Brussels (IB2), VUB-ULB, Brussels, Belgium; Unité de Chronobiologie théorique, Université Libre de Bruxelles (ULB), Brussels, Belgium

## Abstract

Toxin-antitoxin (TA) systems in bacteria and archaea are small genetic elements consisting of the genes coding for an intracellular toxin and an antitoxin that can neutralize this toxin. In various cases, the toxins cleave the mRNA. In this theoretical work we use deterministic and stochastic modeling to explain how toxin-induced cleavage of mRNA in TA systems can lead to excitability, allowing large transient spikes in toxin levels to be triggered. By using a simplified network where secondary complex formation and transcriptional regulation are not included, we show that a two-dimensional, deterministic model captures the origin of such toxin excitations. Moreover, it allows to increase our understanding by examining the dynamics in the phase plane. By systematically comparing the deterministic results with Gillespie simulations we demonstrate that even though the real TA system is intrinsically stochastic, toxin excitations can be accurately described deterministically. A bifurcation analysis of the system shows that the excitable behavior is due to a nearby Hopf bifurcation in the parameter space, where the system becomes oscillatory. The influence of stress is modeled by varying the degradation rate of the antitoxin and the translation rate of the toxin. We find that stress increases the frequency of toxin excitations and decreases the excitation time. The inclusion of secondary complex formation and transcriptional regulation does not fundamentally change the mechanism of toxin excitations. Therefore, the deterministic model used in this work provides a simple and intuitive explanation of toxin excitations in TA systems.

**Author summary**

The genomes of most bacteria and archaea encode several toxin-antitoxin (TA) systems, small genetic elements consisting of the genes coding for an intracellular toxin and an antitoxin that can neutralize this toxin. For several toxins, the target is the mRNA. In this theoretical work we use deterministic and stochastic modeling to explain how toxin-induced cleavage of mRNA in TA systems can trigger large transient spikes in toxin levels. By using a simplified network where secondary complex formation and transcriptional regulation are not included, we show that a two-dimensional, deterministic model captures the origin of such toxin excitations. By systematically comparing our findings with more complex models, we find that the intrinsically stochastic TA system shows similar toxin excitations that are accurately predicted by the underlying simplified theory. This has allowed us to understand how the TA system responds in the presence of stress conditions, finding that toxin excitations become shorter and more frequent.

## Introduction

Toxin-antitoxin modules are small genetic elements, omnipresent on the genomes of bacteria and archaea, that code for a small intracellular toxin and its counteracting antitoxin [1–3]. The antitoxin typically has a higher *in vivo* turnover rate than the toxin [4]. In type II toxin-antitoxin modules, both the toxin and the antitoxin are proteins and the toxin neutralization occurs through the formation of non-toxic complexes [5]. In several toxin-antitoxin modules one antitoxin can neutralize up to two toxins, forming either the complex AT or the complex TAT (see Fig. 1A). Toxin-antitoxin modules further have an intricate transcriptional regulation: the antitoxin has a DNA-binding domain with which it can bind to the promoter/operator region of the toxin-antitoxin module, and functions as a weak repressor. The toxin can function as a corepressor or a derepressor for the antitoxin, depending on the toxin:antitoxin ratio [2]. Different toxins have different targets in the cell, for example, CcdB poisons DNA gyrase [6], while MazF and RelE cleave mRNA [7–9]. Such endoribonuclease toxins will be the focus of this paper.

**Fig 1.**
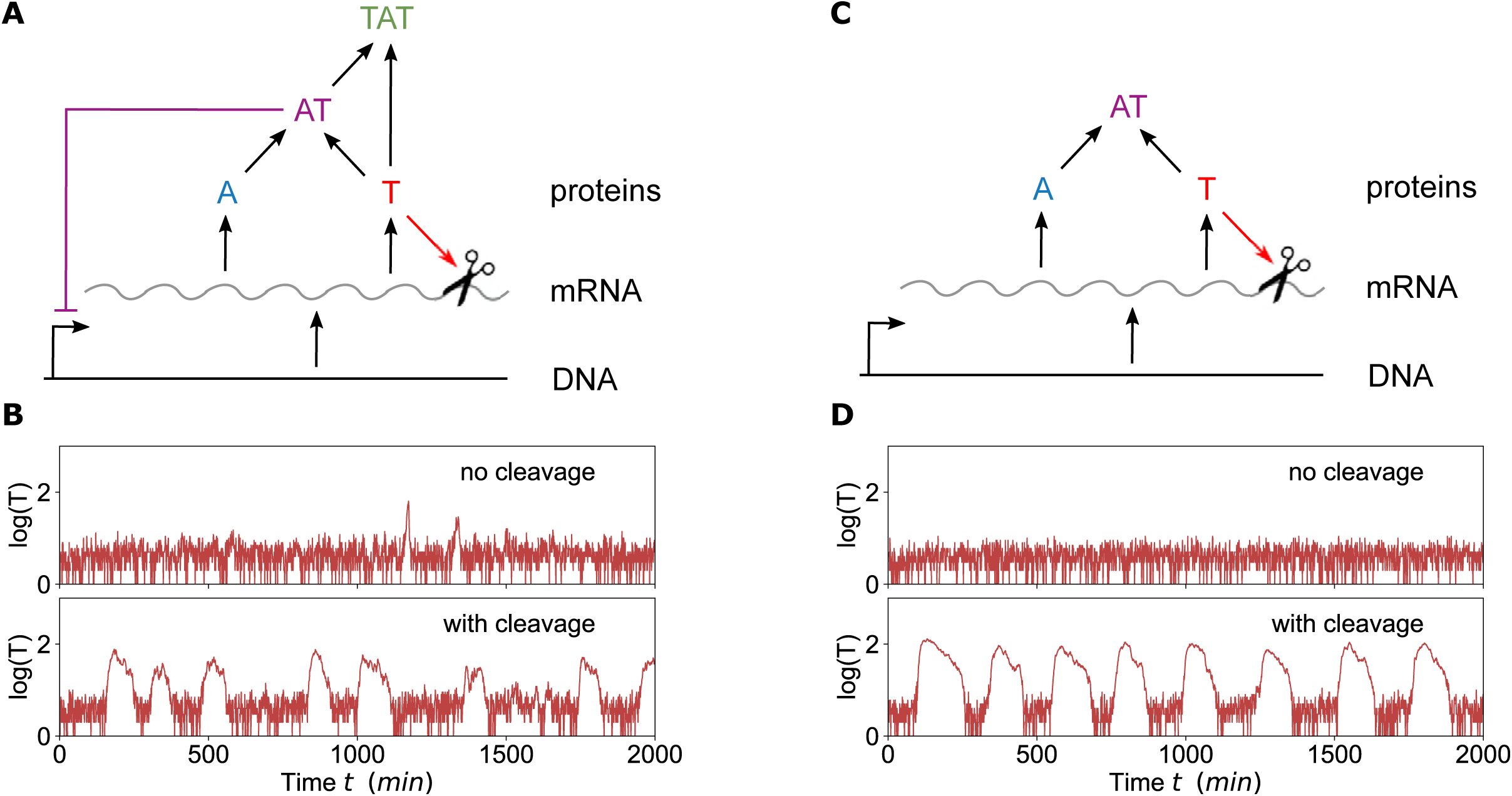
Excitations occur when there is cleavage of mRNA by the toxins. (A) Scheme of the system: DNA is transcribed to mRNA which is translated to toxins T and antitoxins A. These can combine to form the complexes AT and TAT. AT can bind to the DNA and influence the transcription of the mRNA. If the toxin level reaches a threshold this will cause cleavage of the mRNA. (B) Gillespie simulation of the system with TAT and AT binding, with and without mRNA cleavage. The system is only excitable when mRNA cleavage is included. Parameters: *β* = 100, *ε* = 0.074, *γ* = 7.38, *α* = 6.73, *δ*_*c*_ = 1.96, *δ*_*AT*_ = 1.57, *δ*_*a*_ = 10, *κ* = 0.407, *n* = 4. (C) Scheme of the system when formation of TAT and binding of AT are left out, we focus on this system as a minimal model that explains toxin excitations. (D) Gillespie simulation of the system without TAT and AT binding. The same effect as in the complete system is observed: the system is only excitable when mRNA cleavage is included. Parameters: *β* = 100, *ε* = 0.074, *γ* = 7.38, *α* = 6.73, *δ*_*c*_ = 1.96, *δ*_*AT*_ = 1.57, *δ*_*a*_ = 5, *κ* = 0.407, *n* = 4.

Although toxin-antitoxin modules are widespread in prokaryotes, their biological role is currently still unclear. Toxin-antitoxin modules have been implicated in plasmid maintenance, abortive phage infections, the response of bacterial cells to nutritional stress and the formation of persister cells [3, 10]. These are cells that are tolerant to multiple antibiotics because they are in a temporary state of dormancy [11]. Such a period of persistence has been attributed to an increase in the free toxin concentration. Although previously all known type II mRNA endoribonuclease toxins in *E. coli* K-12 were proposed to be involved in persistence, the role of these toxin-antitoxin modules in persister generation in the absence of stress is currently uncertain [3, 12].

Computational studies can be useful to gain insight into the possible dynamics caused by the architecture of the genetic network and the protein-protein, protein-DNA and protein-RNA interactions in a toxin-antitoxin module. Several groups have studied toxin-antitoxin modules computationally, using either deterministic [13–15] or stochastic [16, 17] approaches. From these modeling efforts, two possible deterministic explanations have emerged for the elevated toxin levels that are linked to the generation of persisters. First, it is plausible that there is bistability between a growing, antitoxin-dominated state and a toxin-dominated state linked to persistence [13–15, 17, 18]. A critical component to allow the existence of a toxin-dominated state is that higher toxin levels decrease the cellular growth rates, which in its turn has an effect on the accumulation rate of the toxin itself. Increased noise levels in the presence of stress could lead to stochastic switching between these two states. A second possibility is that the toxin-dominated state only exists as a transient excursion in the free toxin level [16]. Such deterministic excursions could theoretically be generated through a process called excitability, where noise could act to trigger them. Furthermore, if toxins induce growth rate reduction, the duration of such toxin excursions could be significantly lengthened. So far, theoretical studies have only observed such transient toxin excitations using stochastic simulations [16], and a potential link to deterministic excitability remains to be shown. Finally, it is important to note that these different types of deterministic dynamics aim at describing the behavior of single cells. Both bistability and excitability can give rise to bimodal distributions on a population level.

In this article we focus on the effect of the cleavage of mRNA in the presence of elevated toxin levels, which has been recently shown to cause toxin excitations [19]. We use a simplified system, where we leave out the formation of the complex TAT and the transcriptional regulation, as this is the simplest toxin-antitoxin model system that still displays the spikes in the free toxin concentration. We use a deterministic set of differential equations to describe this system and show that it yields similar results as simulations with a Gillespie algorithm. Combining a deterministic approach with a separation of time scales in the system allows to further reduce the problem to a two-dimensional system, which can be visualized and interpreted in the plane. Moreover, it facilitates bifurcation analyses that show how changes in system parameters affect the TA dynamics. For example, we verify how nutritional stress, which causes an increase in the degradation rate of antitoxin, influences free toxin spikes. Finally, we examine how additional feedback mechanisms like transcriptional regulation by binding of the toxin-antitoxin complex to DNA can affect the toxin level.

## Materials and methods

Ordinary differential equations are simulated with the integrate function of the package *scipy* [20] of *python* (Python Software Foundation. Python Language Reference, version 2.7. Available at http://www.python.org). In order to perform bifurcation analysis we use a Newton-Raphson method to solve a set of equations using the jacobian of the system [21]. The maxima and minima of limit cycles are calculated using XPPaut [22].

The stochastic simulations were performed using a Gillespie algorithm, based on treating the biochemical reactions as discrete stochastic events [23], implemented in MATLAB. A detailed description of the different deterministic and Gillespie models used can be found in the supplementary material (S1 File).

## Results

### Toxin-induced mRNA cleavage leads to toxin excitations

Recently, it has been demonstrated experimentally that MazF toxins cleave mRNA in stressful conditions [19], leading to cell growth heterogeneity. Moreover, using stochastic Gillespie modeling, the authors showed that this mechanism induces toxin excitations. This behavior is shown in Fig. 1B using a similar model and parameters as in [19]. Interestingly, we find that toxin excitations are only present if the cleavage of mRNA by the toxins is included in the model (Fig. 1B), suggesting that mRNA cleavage plays an essential role in triggering large spikes in toxin levels. This made us wonder whether other interactions such as transcriptional regulation by binding of AT to the DNA and the formation of the second complex TAT are also required to generate toxin excitations. When removing these interactions altogether, we found that the qualitative behavior of the system was not affected (Fig. 1C-D). With mRNA cleavage, similar toxin spikes were observed, while they disappeared when also abolishing the mRNA cleavage. These simulations showed that toxin-induced mRNA cleavage is the main mechanism by which toxin excitations can be triggered.

### Toxin excitations are the result of underlying deterministic excitability

In order to better understand the dynamical origin of the observed spikes in toxin levels (Fig. 1), we constructed and analyzed a deterministic model of the reduced TA network, shown in Fig. 1C, consisting of ordinary differential equations (ODEs). By not considering the formation of the complex TAT and the transcriptional regulation for now, we focus on analyzing the role of toxin-induced mRNA cleavage in generating toxin excitations. The reduced deterministic model is given by the following 4 ODE equations:

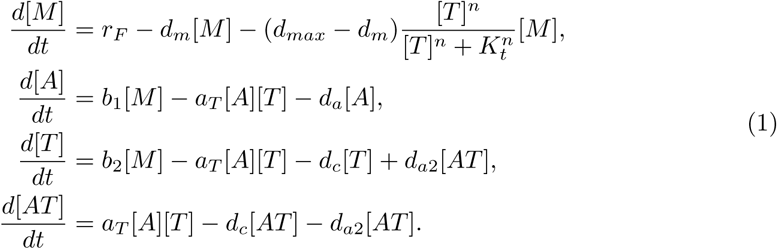

The variables and parameters can be found in table 1 and table 2, where most parameters have been experimentally measured or motivated, see [16, 19]. DNA is transcribed to mRNA [M] with a constant transcription rate *r*_*F*_. This mRNA is then translated to antitoxins and toxins with rates *b*_1_ and *b*_2_. A toxin and an antitoxin can bind to form a complex AT with a rate *a*_*T*_. The mRNA has a degradation rate *d*_*m*_ under normal conditions (in the absence of stress conditions). If the toxin level reaches a threshold *K*_*t*_, then it is assumed there is mRNA cleavage, which is modeled by an increase of the degradation rate to *d*_*max*_ and by using a Hill-function with threshold *K*_*t*_ and coefficient *n*. The antitoxin is degraded with rate *d*_*a*_, whereas the toxin and complex AT are degraded with rate *d*_*c*_. The antitoxin has a faster decay than the toxin, so that the complex AT can break down and release the toxin with a rate *d*_*a*2_. Importantly, the antitoxin is assumed to be less stable than the toxin, as it has a higher degradation rate. Moreover, the translation rate is higher as well, introducing a difference in the time scales of A and T. In order to quantify this, we use *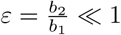*, and reformulate (1) as follows:

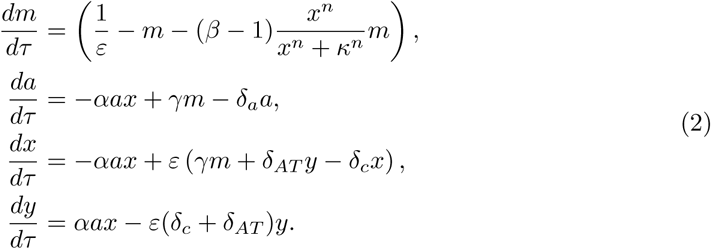

**Table 1.** Normalized variables and their correspondence to the real variables.

| Normalized model |  | Original model |  |
| --- | --- | --- | --- |
| Variable | Definition | Variable | Description |
| $m$ | $\frac{d_m}{r_F \varepsilon} [\text{M}]$ | [M] | mRNA |
| $a$ | $\frac{d_m}{\varepsilon} [\text{A}]$ | [A] | Antitoxin |
| $x$ | $\frac{d_m}{\varepsilon} [\text{T}]$ | [T] | Toxin |
| $y$ | $\frac{d_m}{\varepsilon} [\text{AT}]$ | [AT] | Complex AT |
| $z$ | $\frac{d_m}{\varepsilon} [\text{TAT}]$ | [TAT] | Complex TAT |
| $\tau$ | $d_m \text{t}$ | t | Time |

**Table 2.** Redefinition of the parameters for the normalized model and the correspondence to the original parameters

| Normalized model |  |  | Original model |  |  |
| --- | --- | --- | --- | --- | --- |
| Parameter | Definition | Value | Parameter | Description | Value |
| $\delta_c$ | $\frac{d_c}{d_m \varepsilon}$ | 1.96 | $d_c$ | Degradation rate of T and AT | $0.00028 \text{ s}^{-1}$ |
| $\delta_a$ | $\frac{d_a}{d_m}$ | 1.16 | $d_a$ | Degradation rate A | $0.00231 \text{ s}^{-1}$ |
| $\delta_{AT}$ | $\frac{d_{a2}}{d_m \varepsilon}$ | 1.57 | $d_{a2}$ | Decomposition of AT | $0.000231 \text{ s}^{-1}$ |
| | | | $d_m$ | Degradation of mRNA | $0.002 \text{ s}^{-1}$ |
| $\beta$ | $\frac{d_{max}}{d_m}$ | 100 | $\beta$ | Fold change of mRNA degradation | 100 |
| $\alpha$ | $\varepsilon \frac{a_T}{d_m^2}$ | 6.74 | $a_T$ | Binding of toxin and antitoxin | $0.000365 \text{ M}^{-1} \text{ s}^{-1}$ |
| | | | $r_F$ | Transcription of mRNA | $0.121 \text{ s}^{-1}$ |
| $\gamma$ | $\frac{r_F D}{d_m} b_1$ | 7.38 | $b_1$ | Translation rate of antitoxin | $0.122 \text{ s}^{-1}$ |
| $\varepsilon$ | $\frac{b_1}{b_2}$ | 0.0738 | $\varepsilon$ | Ratio of translation rates | 0.0738 |
| $\kappa$ | $\frac{d_m K_t}{\varepsilon}$ | 0.407 | $K_t$ | Threshold for cleavage of mRNA | 15 M |
| $n$ | | 4 | $n$ | Hill factor | 4 |

This model is now normalized in a way that the parameters are approximately of order *O*(1), except for *β* and *ε*. Here, 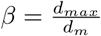 quantifies the difference in mRNA cleavage when it is switched on or off, and *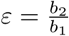* quantifies the difference in the time scales of the system. The definitions of the new variables and parameters are listed in table 1 and table 2. As *ε* is small, three different time scales can be distinguished: of order *O*(*ε*) (slow dynamics), order *O*(1) (fast) and order *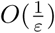* (very fast). The equations for mRNA (*m*) and antitoxin (*a*) are determined by terms of order *O*(1) or order *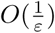* and will approach their equilibrium condition relatively quickly in comparison to the toxin (*x*) and the complex AT (*y*) (*O*(*ε*)). As a result, we assume that 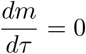 and 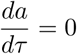 in order to describe the long term dynamics. This is often called a quasi steady state approximation. Carrying out this further simplification the system becomes:

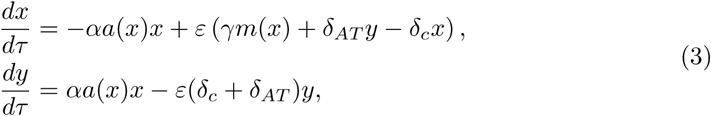

With

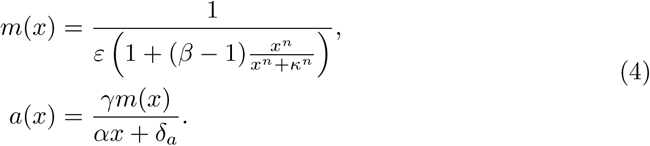

Since this reduced system of equations (3) is two-dimensional, the time evolution of the toxin *x* and complex *y* can be depicted in a simple two-dimensional graph (see Fig 2A). For every set of initial conditions (*x* = *x*_0_, *y* = *y*_0_) in the plane, the equations (3) predict how the system will evolve. In Fig 2, these predictions are illustrated by the gray arrows, which represent the so-called “flow” of the system. The arrows not only show the direction in which the system evolves, but also the speed with which it does so (larger arrows represent faster dynamics). Such a two-dimensional flow allows us to immediately see by eye how the system will evolve in each situation. Another useful tool to understand the behavior of two-dimensional systems is to plot the nullclines. Nullclines consist of all points (*x, y*) where 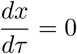 (*NC*(*x*), black line) or 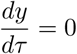 (*NC*(*y*), gray line), as given by:

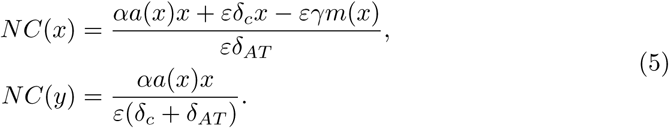

**Fig 2.**
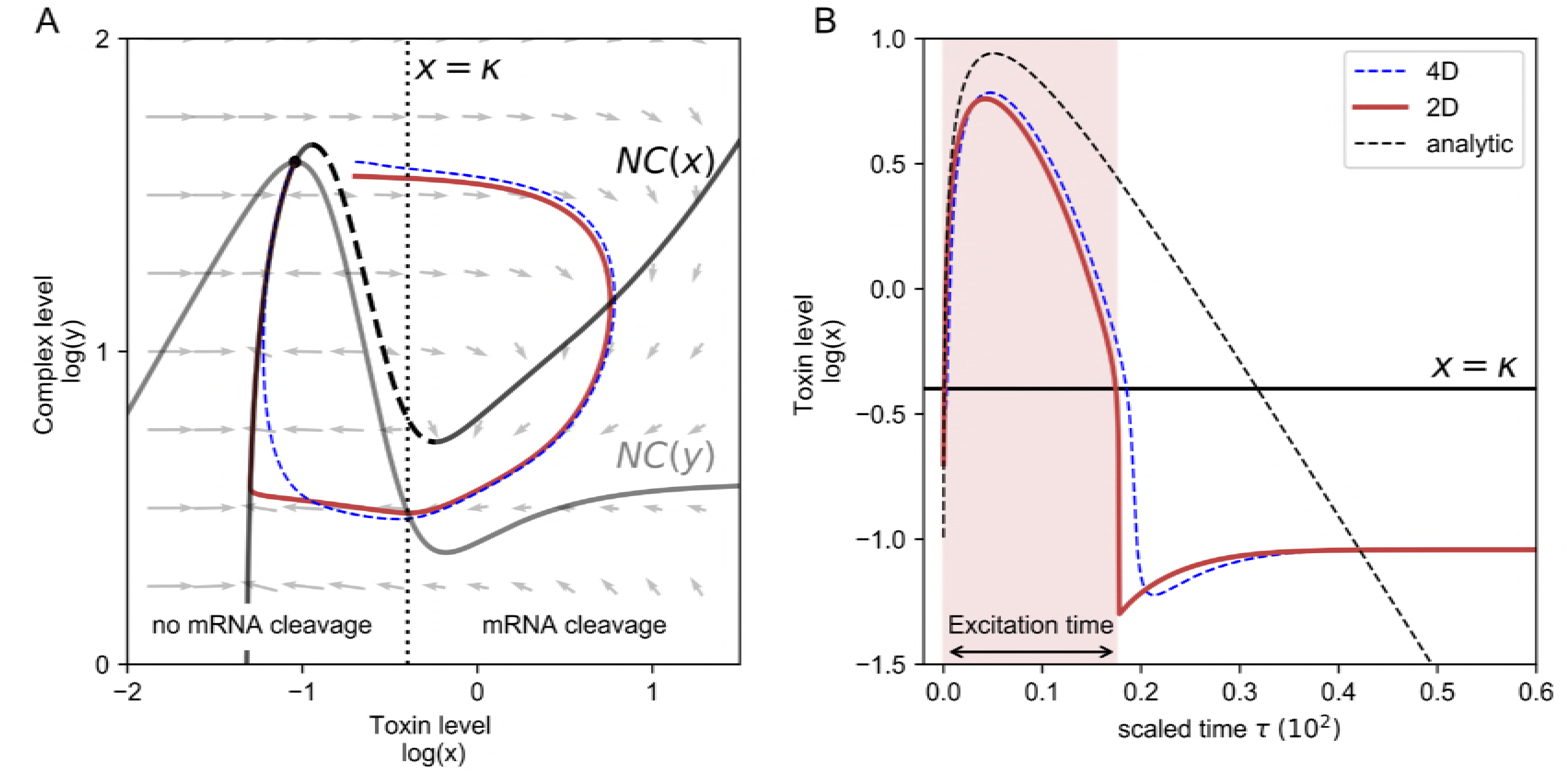
Phase plane visualization (A) and time series (B) of a toxin excitation. (A): Phase plane visualization of the simplified 2D system (3) (red) and the 4D ODE system (2) (blue, dashed). The vectors represent the direction of the flow in each point. NC(x) and NC(y) correspond to the nullclines of the system (2), whereby the excitability threshold is marked by the dashed line. The equilibrium is determined by the cross-section of the nullclines. For toxin levels higher than *x* = *κ* the mRNA cleavage is switched on. In (B) we show the same trajectories as a time course, together with the analytic estimate (6) of the excitation. We define the excitation time as the time that the toxin level is higher than the threshold for the mRNA cleavage, *κ*. (*δ*_*a*_ = 4, *ε* = 0.074)

The intersection of both nullclines represents the equilibrium situation of the system, which is called a fixed point. In our case, there is only one such fixed point and it is stable in the sense that small perturbations will immediately return to this equilibrium. In fact, as it is the only fixed point, all initial conditions (all points in the plane) will eventually return to this equilibrium state. However, when perturbing the system such that the dashed part of NC(x) is crossed, the system will make a large excursion in the plane before returning to the fixed point (see red line in Fig 2A). The excursion in the plane corresponds to a spike in the toxin levels, see Fig 2B. This behavior is called *excitability* [24].

Strictly speaking this analysis in the plane only applies to the reduced two-dimensional system (3). However, we also evaluated the full four dimensional system (2) and projected its dynamics onto the plane (*x, y*) (see blue line in Fig 2A). The full model behaves very similarly, exhibiting similar excitable dynamics. This confirms the validity of the quasi steady state approximation used in deriving the reduced model system. Small differences are observed for low toxin levels, in the regime *x* < *κ*, because the time scale separation used in our reduction no longer applies. In this regime, a different normalization could be used, see supporting information, S2 File.

What happens biologically is that when crossing the threshold for mRNA cleavage (*x > κ*), mRNA cleavage quickly reduces the levels of mRNA (*m*), which in turn suppresses the translation of antitoxin (*a*) and toxin (*x*), thereby preventing the formation of complexes (*y*). Even though toxin translation is suppressed, at first toxin levels keep increasing as the complexes fall apart with a rate *δ*_*AT*_. When there is an insufficient amount of complexes left, the degradation of the toxins, with rate *δ*_*c*_, will dominate. When the level of toxins drops below the mRNA cleavage threshold again, the toxin excitation is terminated, and the system relaxes to the fixed point with low levels of toxins. These rates *δ*_*c*_ and *δ*_*AT*_ determine the shape of the toxin excitation (Fig 2 B) as can also be seen analytically by simplifying the equations (3) in the limit *x >> κ* as follows (for details see supporting information, S2 File):

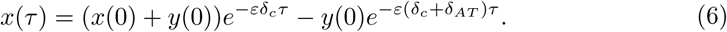

While our findings show that toxin excitations in a deterministic system are related to excitability, it does not yet prove that the toxin spikes we observed in Fig. 1 and in [19] in a stochastic system were indeed related to underlying deterministic excitability. In order to test this, we performed simulations with a Gillespie model, using similar parameter values as for the ODE model in Fig 2. We then visualized the data obtained from the stochastic simulation as a heat-map in the plane (*x, y*), thus plotting the probability to find the system at a certain location in this two-dimensional space. The results are shown in Fig 3A, where the heat-map illustrates the most probable trajectory of the toxin excitation. This most probable path in phase-space nicely overlaps with our deterministic predictions. The corresponding time-traces of the deterministic and the Gillespie simulations are represented in Fig 3 (B) and (C). Together, these simulations confirm that the toxin pulses observed in Gillespie simulations correspond to stochastically triggered pulses, existing due to underlying deterministic excitability.

**Fig 3.**
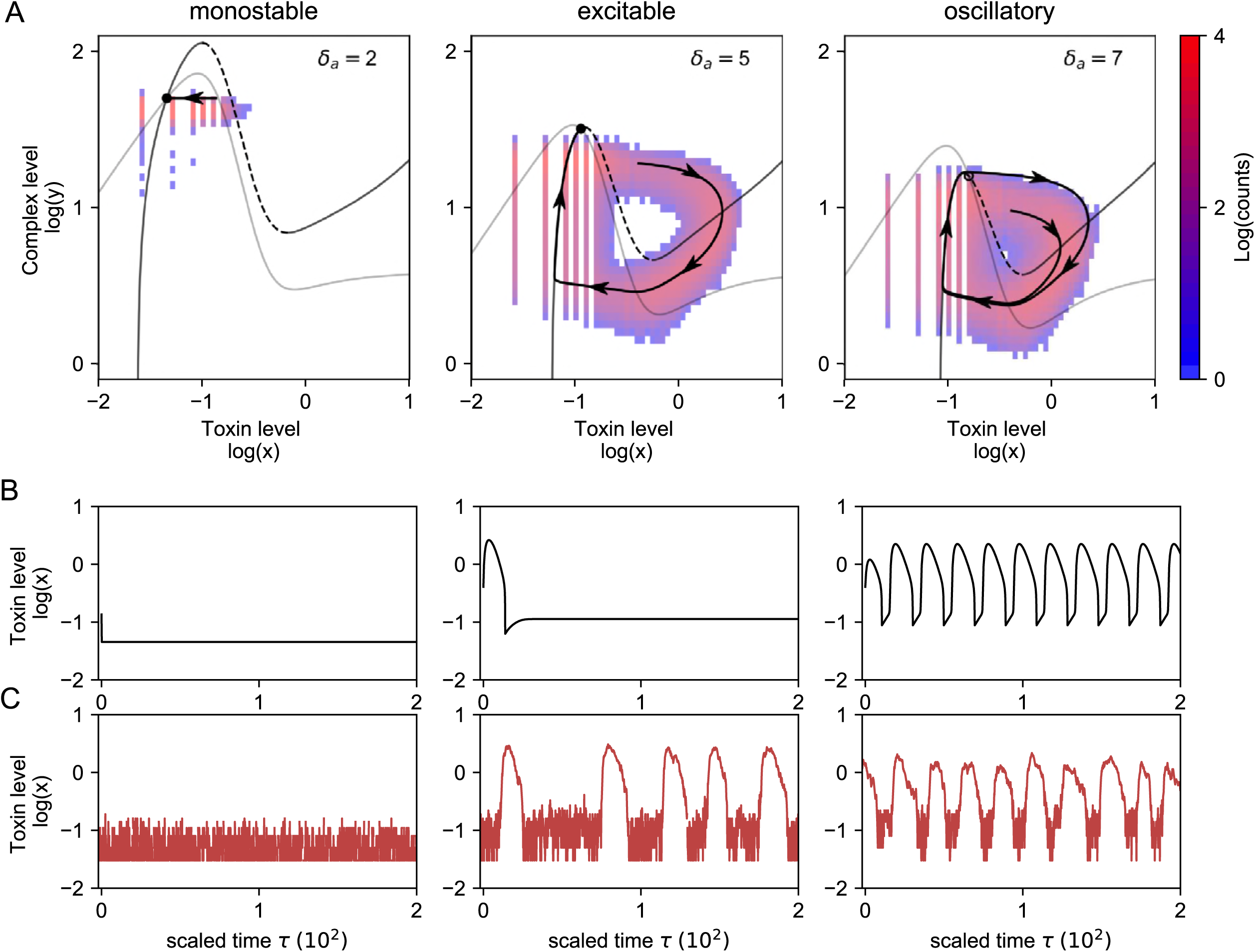
Phase planes (A) and time series of the 2D system (3) (B) and the Gillespie model (C) for monostable, excitable and oscillatory behavior. A: Phase plane representation for monostable (*δ*_*a*_ = 2), excitable (*δ*_*a*_ = 5) and oscillatory (*δ*_*a*_ = 7) behavior. The heatmap corresponds with the amount of times that the Gillespie simulation passes through a local region in phase space. The black trajectory corresponds with the simulation with Eqs. (3) and follows the path of highest probability obtained from the Gillespie data. B: Time traces for the two-dimensional system (3), corresponding to the trajectory in the phase plane. C: Gillespie simulations. The toxin excitations have similar shape as when simulated with the deterministic models.

### The toxin-antitoxin system can be monostable, excitable or oscillatory

Next, we explored the robustness of these toxin excitations to changes in the system parameters. In Fig 3, we show that the system can show qualitatively different behavior when changing the antitoxin degradation rate (*δ*_*a*_). For *δ*_*a*_ = 2, the system has one stable solution and stochastic Gillespie simulations do not show any toxin excitations. This is explained by the fact that the excitation threshold is too large, such that noisy excursions around the stable state are too small to trigger any excitation. We note that the projected Gillespie data does seem to occasionally cross the threshold (dashed line), even though this does not lead to an excitation. This discrepancy between the Gillespie simulations and the reduced 2D ODE system is explained by the fact that perturbed values of *m* and *a* in the Gillespie simulation do not immediately reach their quasi steady state condition. Whereas an excess of toxins immediately leads to mRNA cleavage in the reduced deterministic case, there is a small delay in the Gillespie simulations. As long as the antitoxin level is sufficiently large, complex formation is able to reduce the toxin level, thus having a stabilizing effect on the system.

When increasing *δ*_*a*_, the excitability threshold is reduced, such that the probability to stochastically excite toxin spikes increases. Indeed, for *δ*_*a*_ = 5, toxin excitations can be observed in the Gillespie simulations. Although the deterministic 2D model accurately predicts the shape of the toxin excitations, the equilibrium state is never fully reached in the Gillespie simulations. The reason is that the fixed point is situated close to the top of the nullclines, where the excitability threshold is significantly smaller than the noise. Therefore, the system will never relax to its fixed point and instead a new excitation will be triggered. This discrepancy disappears for increasing excitability threshold (see supporting information, S3 Fig).

Finally, for even larger antitoxin degradation rates (e.g. *δ*_*a*_ = 7), the reduced ODE system shows oscillatory behavior. The fixed point becomes unstable and instead the system converges to a limit cycle. Consistent with these deterministic findings, Gillespie simulations also show more regular excitations. The deterministic prediction of the limit cycle and the most probable path as given by the heat-map are somewhat shifted with respect to one another. Similarly as in the excitable case, noise will cause the system to cross the threshold *NC*(*x*) (dashed line) earlier than in the deterministic case. When the initial condition is such that it corresponds with a point that is situated in the main band of the Gillespie data, then the deterministic trajectory of the first excitation does correspond well with the Gillespie simulations.

### Cellular stress causes more, but shorter, excitations

When a bacterial cell is experiencing nutritional stress such as amino acid starvation, the degradation rate of several antitoxins (*δ*_*a*_) increases due to the increased activity of cellular proteases such as Lon [7, 25]. As shown before in Fig 3, this increases the probability of toxin excitations. Another way to influence the toxin level is by directly increasing the translation rate of the toxin (*ε*). Here, using time evolution simulations and bifurcation analysis, we analyze how changes in the system parameters *δ*_*a*_ and *ε* affect the toxin-antitoxin dynamics. Other important biological parameters that affect the dynamics are the parameters determining the mRNA cleavage (*β, n, κ*) and the binding parameter *α*. The influence of these parameters on the dynamics and shape of the nullclines are discussed in S1 Fig.

In order to analyze the results of the bifurcation analysis (Fig 4) we define the excitation time as the time that the toxin level is higher than the toxin threshold *κ* used in the Hill function (see Fig 2 (B)). Fig 4(A) and (B) show the average excitation time of toxin excitations, obtained with deterministic (A) and stochastic Gillespie (B) modeling, as a function of the antitoxin decay rate *δ*_*a*_ and toxin translation rate *ε*. Our simulations show that excitations occur for moderate stress levels, while higher stress leads to oscillatory behavior. When stress levels are increased even further, those oscillations eventually disappear as the fixed point becomes stable again. This loss of oscillations is less clear in the Gillespie data, as some excitations are still detected. This is explained by the fact that the toxin value *x* at the fixed point lies close to threshold *κ*, such that noise can still trigger excursions. Interestingly, the excitation time is larger for lower values of *δ*_*a*_ and *ε*. The reason is that here the fixed point is situated closer to the top of the nullclines, so that the amplitude of an excitation is larger.

**Fig 4.**
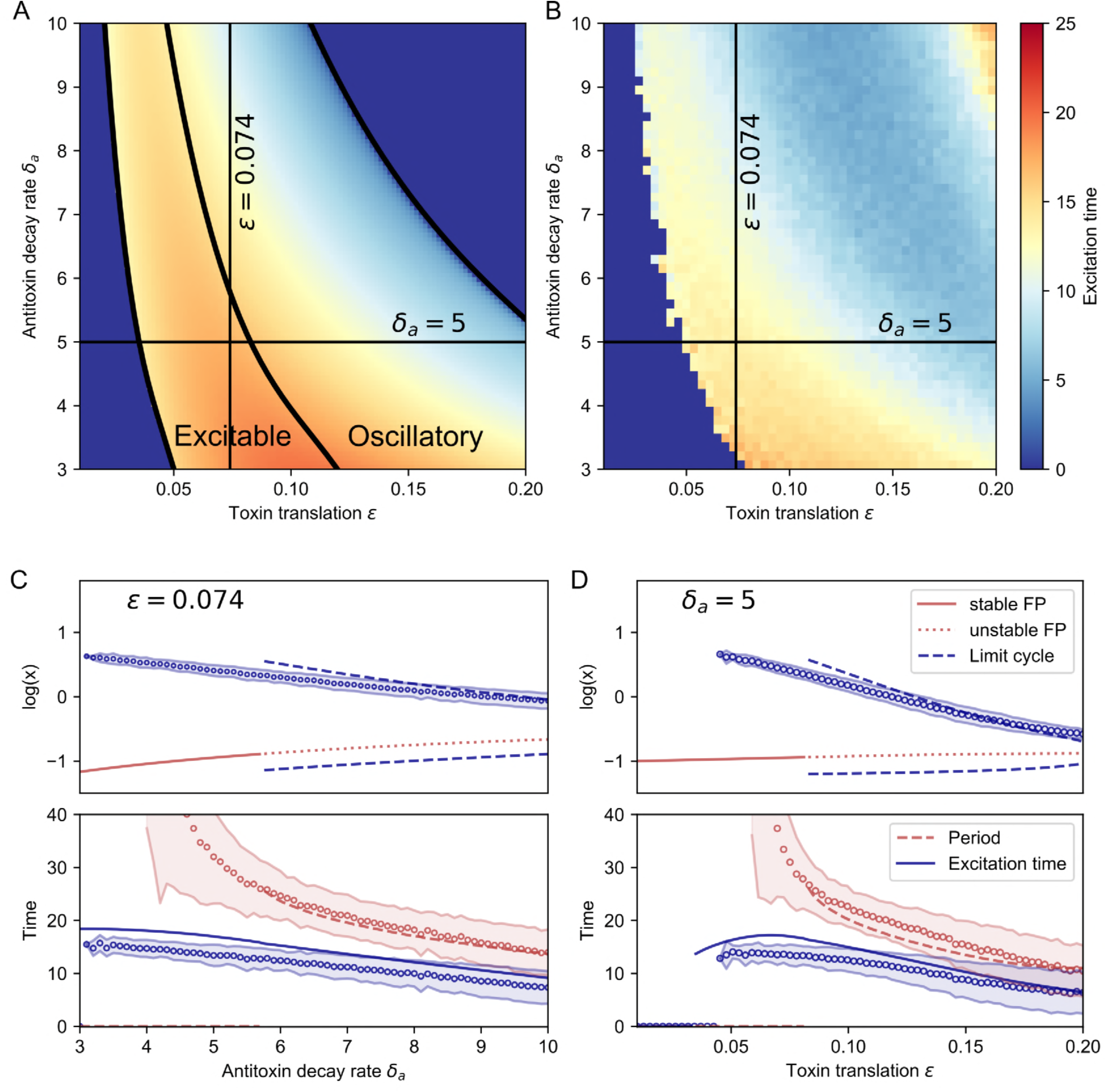
Influence of the antitoxin decay rate *δ*_*a*_ and the toxin translation rate *ε* on the excitation time. A. Heatmap of the excitation time as a function of the antitoxin decay rate *δ*_*a*_ and the toxin translation rate *ε*, using Eqs.(3). B. Heatmap of the excitation time as a function of the antitoxin decay rate *δ*_*a*_ and the toxin translation rate *ε*, using Gillespie simulations. C. Bifurcation diagram for the antitoxin decay rate *δ*_*a*_, keeping *ε* = 0.074 constant. The system becomes oscillatory due to a subcritical Hopf bifurcation. The period and excitation time in the deterministic (solid, dashed and dotted lines) and the Gillespie (mean: open circles, shaded region: mean *±* one standard deviation) model are compared and show a good correspondence. D. Bifurcation diagram for the toxin translation rate *ε*, keeping *δ*_*a*_ = 5 constant. Notation same as in C.

This is analyzed in more detail by keeping *ε* = 0.074 fixed in Fig 4(C) and by keeping *δ*_*a*_ = 5 fixed in Fig 4(D). The bifurcation analysis, using Eq.(3), shows that the fixed point loses its stability in a Hopf bifurcation. We found that the amplitude of the limit cycle is increased instantaneously, as this is a subcritical Hopf bifurcation. This means that the fixed point merges with an unstable limit cycle (not shown), thereby losing its stability. The excitability of this system is related to the vicinity of a limit cycle in parameter space. As the period is non-diverging at the bifurcation point, this is an example of type II excitability [26]. The maximum toxin level during an oscillation, the excitation time and the period of an oscillation are compared with the observation in the Gillespie data and show a good correspondence. However, the maximum toxin level and the excitation time show a discrepancy, due to the fact that the Gillespie data does not fully reach the fixed point, as explained in last section. This behavior is illustrated in more detail in S2 Fig, where we explore the system dynamics for different values of *ε*. In Fig 4(C) and (D) we clearly see that the period decreases for increased stress, so that excitations are more frequent. However, the excitation time decreases as well, although not as significantly as the period. Therefore, for high amounts of stress, there could be more cells in the toxic state with a quicker return to the normal state.

### Toxin excitations persist when including additional complex formation and DNA binding

So far, we showed that toxin-induced mRNA cleavage is the main mechanism leading to excitable behavior. Here, we will incorporate the effect of a second complex TAT and the binding of AT to the DNA (see Fig. 1A) and show that this does not change the dynamics in a qualitative manner. Each deterministic and its corresponding stochastic model is described in more detail in S1 File.

#### Inclusion of a secondary complex

The system (3) can be extended to incorporate the second complex TAT by using an additional variable *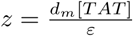*. As a simplification we assume that *y* and *z* have the same creation and decay rates *α, δ*_*c*_. We also assume that *z* breaks down with rate *δ*_*AT*_ into two toxins *x* and one antitoxin *a*. The difference in time scales still exists, so that *m* and *a* can be assumed to be in steady state, and the resulting normalized equations are the following:

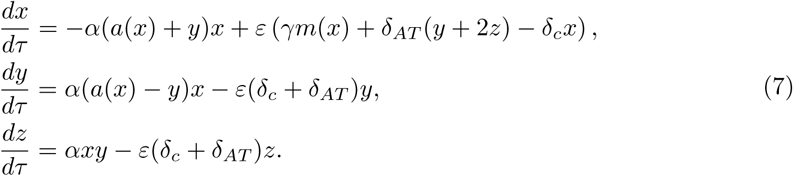

To show the similarity of this three-dimensional ODE model with the two-dimensional model (3), we look at the total amount of toxin in the complexes, *c* = *y* + 2*z* rather than the dynamics of *y* and *z* separately, resulting in the following system:

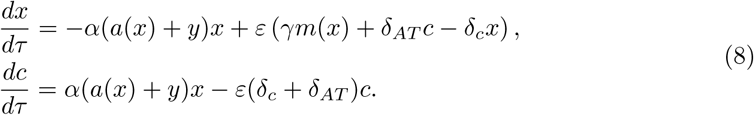

The only difference between these equations and (3) are the terms of *α*(*a* + *y*)*x*. However, during an excitation the terms of order **O**(*ε*) become important, which have the same shape as in (3). Hence, the mechanism of the excitation does not change by inclusion of the secondary complex *z*. The altered term −*α*(*a* + *y*)*x* does have a stabilizing effect on the system dynamics, as *x* will be degraded more quickly. This causes a shift in the nullclines such that this system requires a larger perturbation to excite toxin spikes. This is shown in Fig 5, where the plotted nullclines are approximations, assuming that *a* and *y* are of the same order of magnitude. This way the terms *α*(*a* + *y*)*x* reduce to 2*αax*. As a result there is not an exact correspondence with the intersection of the nullclines and the fixed point of the system.

**Fig 5.**
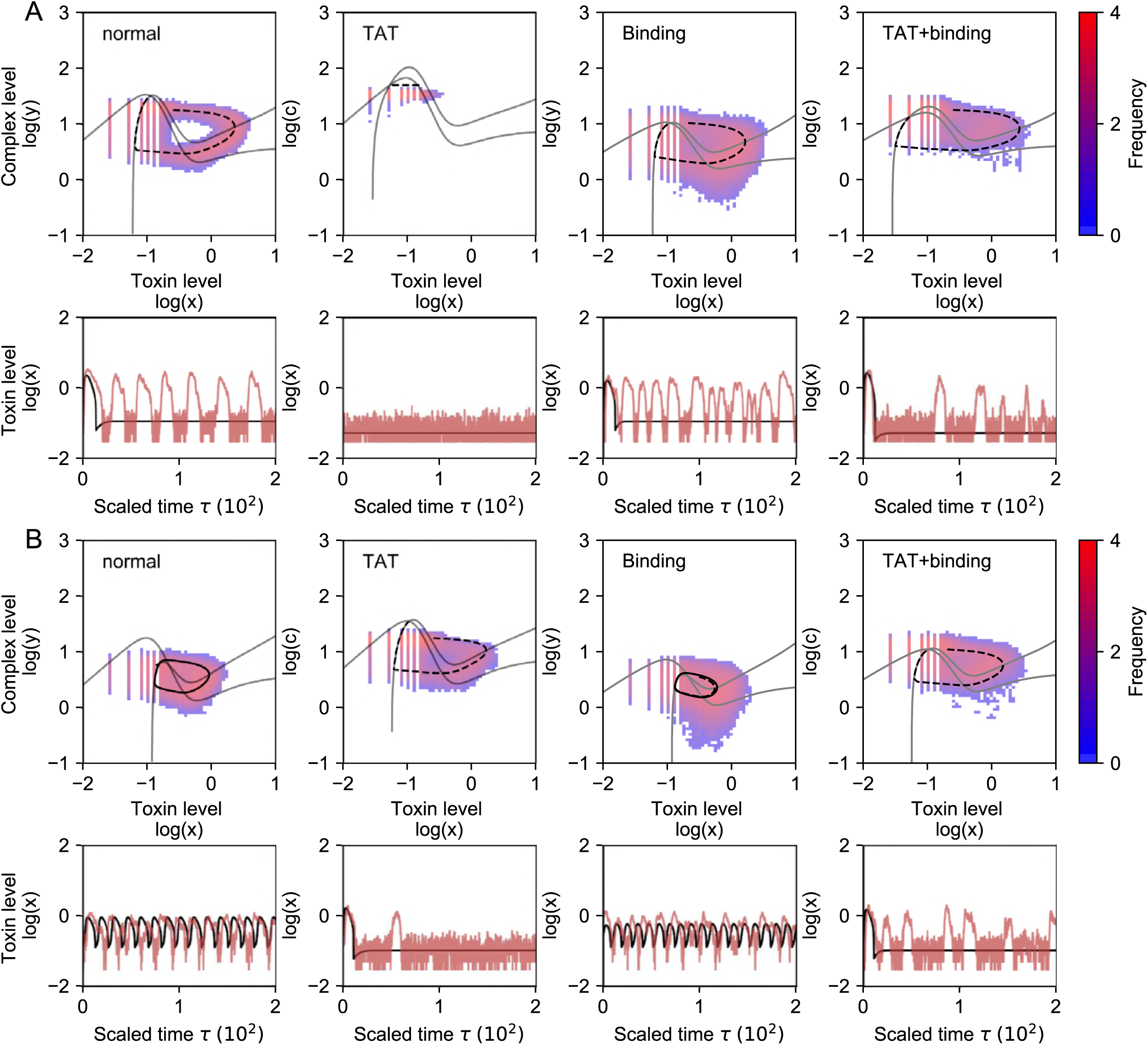
Inclusion of secondary complex formation and trancription regulation by DNA binding does not change the qualitative dynamics. In the simulations with DNA binding we used *r* = 4.9 as binding parameter, see S1 File. A: Simulations for the different models using *δ*_*a*_ = 5. The inclusion of the secondary complex TAT stabilizes the system, no excitations are observed in the Gillespie data. The inclusion of binding of AT to the DNA destabilizes the system as the frequency of excitations increases. When both mechanisms are included, the frequency of the excitations is reduced. B: Simulations for the different models using *δ*_*a*_ = 10. By artificially increasing the amount of stress the system with the secondary complex TAT becomes excitable as well. The system with DNA binding is oscillatory. The system with TAT as well as binding shows excitable dynamics.

#### Inclusion of binding to DNA

Transcriptional regulation occurs when the complex AT (*y*) binds to the DNA and thereby inhibits the transcription of mRNA. As such, the translation of the toxins and antitoxins is also indirectly inhibited. In the deterministic system, we model DNA binding by incorporating an inhibiting Hill function *r/*(*r* + *y*) into the translation rate (for derivation, see supporting information, S1 File). This expression decreases from 1 (no suppression) to 0 (complete suppression) for increasing values of the complex *y*. When assuming quasi steady state conditions for *m* and *a*, the system is still described by (3), but the expression for *m*(*x*) is changed as follows:

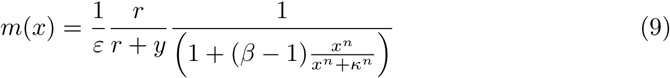

As the binding inhibits the translation of toxin and antitoxin, there is less complex formation. This causes a shift in the nullclines leading to an equilibrium with less complexes (Fig 5). Binding of AT to the DNA did not stabilize the system as the threshold for toxin excitations did not increase. Therefore we found a comparable frequency of excitations as in the absence of DNA binding. Even though there is no stabillizing effect, the advantage of DNA binding is that the cell uses less energy as there is less translation [16]. When both the secondary complex TAT and DNA binding are included in the system (Fig 5), then there are less complexes due to the binding and the system is more stable due to TAT.

### Conclusions

TA modules are small dynamic systems, coding for a toxin and its corresponding antitoxin [1–3]. The toxin level can affect a cell in different ways: post-segregational killing, abortive phage infections and the formation of persister cells [3, 10], although the latter is currently unclear [3, 12]. There exist different dynamic explanations for elevated toxin levels in cells. The first possibility is that this corresponds to a second equilibrium state, due to bistable behavior [14, 15, 17, 18], while the second possibility is that stochastic perturbations cause pulses in the toxin level, corresponding to excitable behavior [16]. Recently it was found that mRNA cleavage by the toxins can cause toxin excitations, leading to cell growth heterogeneity [19].

In this article we used a deterministic model to analyze the exact mechanism behind such excitations. As there is a difference in time scales between toxin and antitoxin translation and degradation, the model can be simplified to a set of two ordinary differential equations (ODEs), allowing a description in the phase plane. An excitation occurs when a threshold is crossed, mRNA cleavage is switched on, and the repression of translation prevents an immediate return to the equilibrium state. By systematically comparing with Gillespie simulations we showed that even though the real system is inherently stochastic, a deterministic model is capable to describe the observed dynamics. Moreover, in the deterministic system it can be shown that the excitable behavior is due to the vicinity of a Hopf bifurcation where a limit cycle is created. Stress can be modeled by varying the antitoxin degradation rate and the toxin translation rate, which increases the probability of excitations, as the fixed point gets closer to the unstable branch in the phase plane. In conclusion, even though this system is inherently stochastic, we provided a deterministic description of the excitable behavior in TA modules due the presence of toxin-induced cleavage of mRNA.

Similar excitable behavior in bacteria was theoretically and experimentally analyzed in the ComK - ComS gene regulatory circuit in *Bacillus subtilis*, where excitability led cells to be in a transient state in which they were competent to take up DNA from the environment [27, 28]. Although the circuitry of interacting genes and proteins in the ComK - ComS system is significantly different than that of the TA systems we studied here, the type of excitable behavior is similarly caused by a combination of fast positive and slow negative feedback loops. By using quantitative fluorescence time-lapse microscopy to observe circuit components in individual cells, and comparing such measurements with mathematical models, significant new insights were gained into how the ComK - ComS gene regulatory circuit works [27, 28]. We hope that our model will trigger new experimental efforts in the field of TA systems, especially to measure the dynamics of circuit components on a single cell level, and that they will help in shedding new light on the temporal dynamics of cellular toxin levels and growth rates. Such efforts would also allow to bridge the internal dynamics in individual cells and the dynamics on the level of whole cell populations, where bimodal distributions of fast and slowly dividing cells have been observed [19].

## Supporting information

**S1 Fig. Influence of the parameter** *n*, *β*, *κ* **and** *α* **on the phase plane.** These parameters can change the shapes of the nullclines: an increase in *n* makes the slope steaper, an increase of *β* leads to a bigger difference between the nullclines in the toxic and normal state, *κ* changes the threshold itself and *α* affects the complex level at steady state. However, the overall behavior as explained in the main text remains the same.

**S2 Fig. Influence of an increase of** *ε***. Comparison between the deterministic and the Gillespie simulations.** An increase in stress by varying *ε* leads subsequently to excitability and oscilatory behavior. The amplitude of the oscillations decrease with an increase of *ε*, while the fixed point becomes closer to the boundary where *x* = *κ*.

**S3 Fig. Gillespie simulations for small values of** *ε*. The Gillespie simulations do not reach the fixed point when this is situated near the top of the nullclines (*ε* = 0.6 and *ε* = 0.8), as an excitation will occur as soon as the unstable branch is crossed. The data converges to the fixed point more closely when this is not situated in the top of the nullclines (e.g. *ε* = 1.0).

**S1 File. Model descriptions** A description of the different deterministic and corresponding Gillespie models that are used throughout the main text.

**S2 File. Analytic results of the TA system.** Description how to find an analytic solution for the excitation and a discussion about the behavior of the system for *x ≪ κ*.

## Acknowledgements

We thank N. Nikolic, J. Danckaert and D. Gonze, together with members of the Gelens laboratory, for thoughtful comments and discussions.

